# The helicase domain of Polθ counteracts RPA to promote alt-NHEJ

**DOI:** 10.1101/176271

**Authors:** Pedro A. Mateos-Gomez, Tatiana Kent, Ekaterina Kashkina, Richard T Pomerantz, Agnel Sfeir

**Author notes:** Correspondence: Agnel Sfeir Skirball Institute/NYU Langone Medical center 540 First Avenue 4^th^ floor/ Lab3 New York, NY 10016.

## Abstract

Mammalian polymerase theta (Polθ; encoded by *POLQ*) is a unique multifunctional enzyme that promotes error-prone DNA repair by alternative-NHEJ (alt-NHEJ). Here we perform structure-function analyses and report that, in addition to the polymerase domain, the helicase activity plays a central role during Polθ–mediated double-stranded break (DSB) repair. Our results show that Polθ–helicase promotes chromosomal translocations by alt-NHEJ in mouse embryonic stem cells. In addition, the helicase activity suppresses CRISPR/Cas9 mediated gene targeting by homologous recombination (HR). *In vitro* experiments reveal that Polθ–helicase displaces RPA to facilitate the annealing of complementary DNA during alt-NHEJ. Consistent with an antagonistic role for RPA during alt-NHEJ, we show that the inhibition of RPA1 subunit enhances end-joining and suppresses recombination at telomeres. Taken together, our results reveal that the balance between HR and alt-NHEJ is regulated by opposing activities of Polθ and RPA, providing critical insight into the mechanism that control DSB repair pathway choice in mammalian cells.

## Introduction

Double strand break (DSB) repair is essential for cellular survival and maintenance of genome integrity. In addition to the two canonical DSB repair pathways of homologous recombination (HR) and classical non-homologous end-joining (C-NHEJ), an error-prone and mechanistically distinct pathway termed alternative-NHEJ (alt-NHEJ) covalently joins DNA ends^1^. Alt-NHEJ was initially thought to act as a back-up mechanism^2^, however, recent studies indicated that it operates even when HR and C-NHEJ are intact^3^, and is a major repair pathway during early vertebrate development^4^. Moreover, when HR and C-NHEJ are impaired, mammalian cells become highly dependent on alt-NHEJ for survival^5-7^. Whether alt-NHEJ comprises one or multiple overlapping mechanisms is still a matter of debate, yet a significant fraction of its events are characterized by the presence of microhomology, as well as deletions and insertions that scar DNA repair sites^1^. Whole genome sequence analysis of BRCA1/2 mutated tumors identified a genomic signatures with features (insertions and microhomology) that are highly reminiscent of alt-NHEJ repair^8-10^. The source of mutagenic insertions during alt-NHEJ has been attributed to the activity of polymerase theta (Polθ – encoded by *POLQ*), a unique enzyme found only in metazoans^11^.

Polθ was originally identified in *D. melanogaster* through the analysis of mus308 mutants that displayed hypersensitivity to agents that induce DNA interstrand cross-links (ICL)^12^. The activity of Polθ was first linked to alt-NHEJ during P-element transposition in flies^13^, and later found to promote end-joining in response to different DSB-inducing agents in plants^14^, worms^15^, fish^4^, and mammals^5,6,16-18^. Mammalian Polθ stimulates alt-NHEJ in response to endonuclease-cleaved reporter constructs^5,6,16,17^, drives the fusion of dysfunctional telomeres^5^, and promotes chromosomal translocations in mouse embryonic stem cells^5^. In addition to promoting alt-NHEJ, recent evidences suggested that Polθ negatively regulates HR^5,17^. The accumulation of IR-induced Rad51 foci was significantly enhanced upon suppression of Polθ^5,6^. In addition, increased recombination in response to Polθ inhibition was noted at dysfunctional telomeres and using I-SceI-based reporter assays^5,6^.

Polθ is a multifunctional enzyme that is composed of a superfamily 2 (SF2) Hel308-type helicase domain at the N-terminus, a low fidelity A-family polymerase domain at the C-terminus, and a non-structured central domain^11^. The role of the polymerase domain (Polθ–polymerase) during alt-NHEJ was elucidated through a series of biochemical studies^5,17,19-23^. Polθ–polymerase is capable of oscillating between templated and non-templated activities to promote alt-NHEJ. Templated nucleotides are primarily copied from regions flanking the break sites in *cis* and in *trans*, while non-templated extensions by Polθ– polymerase are driven by its template-independent terminal transferase activity^17,19^. The biochemical and biological functions of the helicase domain (Polθ–helicase), which is similar in sequence to Hel308 and RecQ type helicases, remain poorly understood. Analysis of the crystal structure revealed that the helicase domain of Polθ forms a tetramer, an arrangement that was also observed in solution^24^. It has been proposed that the helicase domain acts together with the Rad51 interaction motif within the central region to antagonize strand invasion during HR^6^. Analysis of alt-NHEJ in *D. melanogaster* indicated that cells harboring mutations in the ATPase motif display less microhomology at repair junctions, albeit the overall efficiency of end-joining was not affected^13^. Similar results were obtained upon studying the repair of breaks incurred on episomal end-joining substrates in mouse cells overexpressing inactive Polθ–helicase^7^. So far, biochemical analysis failed to show any DNA unwinding activities, yet its Polθ–ATPase activity is readily detectable and stimulated by DNA^23,25^. Despite that presence of the helicase at the N-terminus of *POLQ* s conserved among metazoans, the mechanism by which it influences DSB repair remains unknown.

How cells choose between erroneous alt-NHEJ and accurate HR is critical, yet poorly understood. Both pathways are maximally active in S and G2 phases of the cell cycle, with alt-NHEJ processing 10-20% of breaks incurred during S phase in HeLa cells^3^. As in HR, the initial step of alt-NHEJ involves DNA end-resection by the Mre11–Rad50-Nbs1 (MRN) complex and CtIP to create short single-stranded DNA (ssDNA) overhangs^3,26,27^. Resection exposes microhomology internal to break sites, which could facilitate spontaneous annealing of ssDNA to form a synapsed intermediate that is essential for end-joining. The stability of the synapsis is governed by the extent of microhomology and influenced by factors acting at break sites. For example, in *S. cerevisiae*, where the frequency of alt-NHEJ activity is very low, end-joining requires 5-12 base-pairs (bp) of homology^28,29,30^. Furthermore, the binding of the replication protein A (RPA) complex to resected ends prevents spontaneous annealing between microhomologous overhangs, thereby hindering alt-NHEJ in yeast^31^. Whether the function of RPA as a negative regulator of alt-NHEJ is conserved in mammalian cells remains unknown. Furthermore, the factors and mechanisms that promote and stabilize annealed intermediates with little microhomology (i.e. ∼2 bp)^32^, typical of mammalian alt-NHEJ, have yet to be determined.

In this study, we identify a crucial role for Polθ–helicase in promoting alt-NHEJ and counteracting HR. Our data demonstrate that the function of Polθ–helicase is consistent with an ATP-dependent annealing helicase that counteracts RPA to drive alt-NHEJ. Lastly, we show that inactivation of RPA subunit 1 enhances alt-NHEJ at the expense of HR. Altogether, our results indicate that opposing activities of RPA and Polθ–helicase at DSBs regulate the balance between alt-NHEJ and HR. As such, the outcome of this interplay will have profound impact on the stability of mammalian genomes and the proliferation of HR-deficient cancer cells.

## Results

### Mutational analysis uncovers the function of independent domains within Polθ

Polθ is the only known eukaryotic DNA polymerase that contains an intrinsic helicase domain, which lies at the N-terminus^23^. Three putative Rad51 interaction motifs were identified within the unstructured central domain of human Polθ, however, only one motif is evolutionary conserved. The function of the different Polθ domains in mammalian cells has been primarily investigated using cellular-based assays that involve overexpression of various truncated alleles of Polθ, such as the helicase domain^6,7,16^. The cellular levels of Polθ in cells are low^33^ and *POLQ* overexpression altered DNA replication and triggered a DNA damage response^34^. As such, it is imperative to address the function of its different domains in the context of physiological Polθ levels. To that end, we exploited CRISPR/Cas9 gene editing and targeted the endogenous *Polq* locus in mouse embryonic stem (mES) cells. We generated independent cell lines with inactivating mutations in the conserved ATPase (K120G)^7^ and polymerase (D2494P, E2495R)^35^ domains (Figure 1a). In addition, we engineered mouse ES cells harboring a 47 aa deletion (ΔD844-M890) that eliminated the conserved Rad51 binding site (Figure 1a). For each mutant, we isolated multiple clonally-derived cell lines and confirmed successful bi-allelic targeting by Sanger sequencing (Figure S1a).

**Figure 1.**
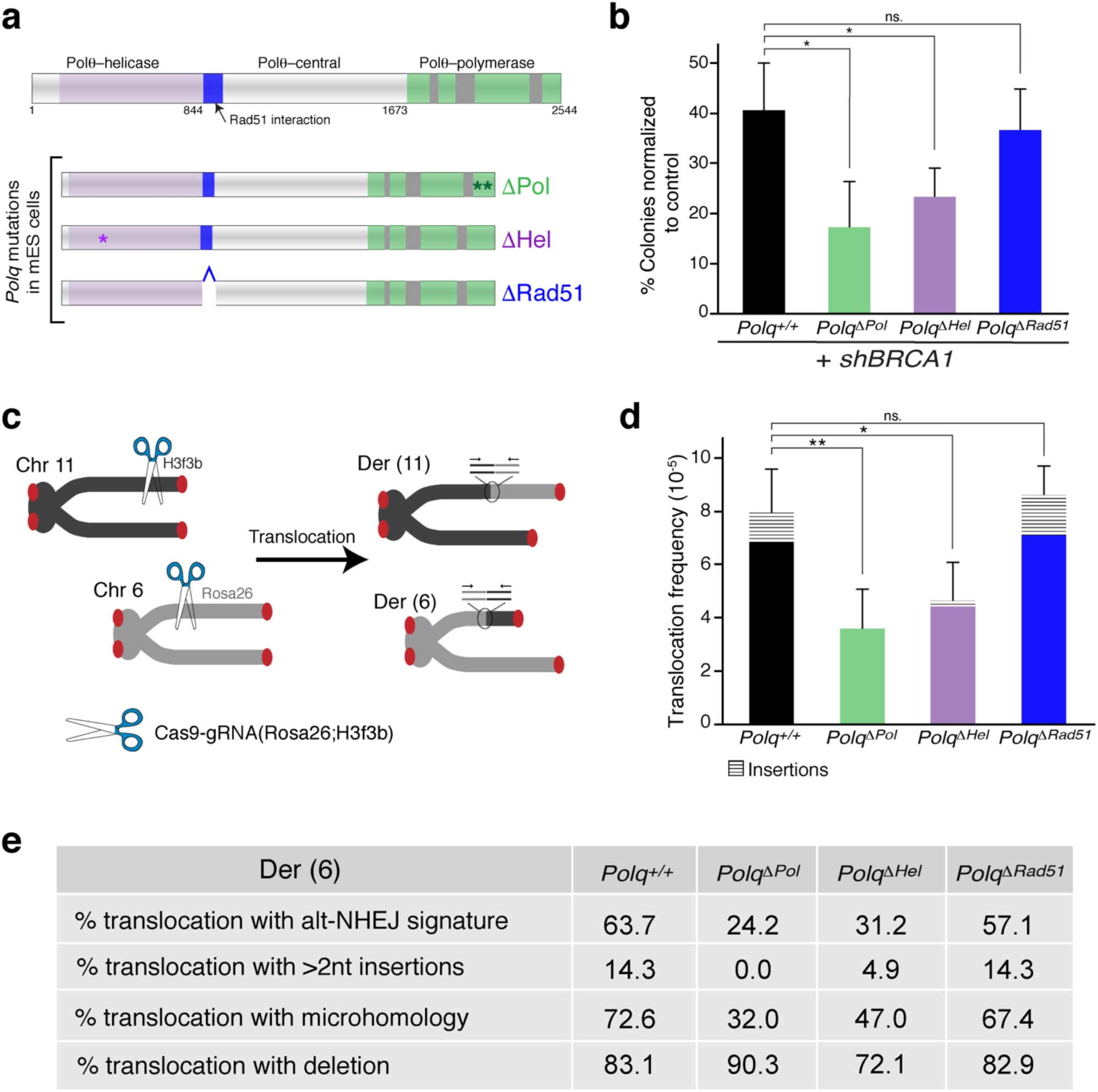
Structure-function analysis reveals the function of Polθ–helicase during alt-NHEJ. **(a)** Top – schematic representation depicting the different domains of Polθ. Three Rad51 binding domains were identified in human *POLQ*, however, only one motif is conserved in mouse (highlighted in blue). The polymerase domain contains three conserved unique insertion loops (highlighted in grey) that are involved in DNA synthesis. Bottom – CRISPR/Cas9 gene targeting was carried out in mouse ES cells (CCE) to introduce two base substitutions (Asp2494 and Glu2495) that inactivate the polymerase domain. In addition, independent gene editing was carried out to inactivate the ATPase activity (Lys120) and delete the conserved Rad51 interaction motif (residues 844-890). (b) Quantitative analysis of colony formation assay in mES cells carrying the indicated *Polq* mutant alleles and treated with sh*BRCA1*. In each cell line, colonies were normalized to cells treated with shControl. Bars represent means of three independent experiments ±S.D. For every experiment, two independent clonally derived cells lines were analyzed. * p<0.05; two-tailed Student’s t-test. **(c)** Schematic of chromosomal translocation assay in which DSBs are induced at the Rosa26 and H3f3b loci. Generation of derivative chromosomes Der(6) is detected by nested PCR. **(d)** Frequency of chromosomal translocation in *Polq*^+/+^, *Polq*^*ΔPol*^, *Polq*^*ΔHel*^, and *Polq*^*ΔRad51*^ cells. Bars represent mean ±S.D. Two independent experiments, each was carried out using two clonally-derived cells lines. *p<0.05; two-tailed Student’s t-test. **(e)** Table summarizing the analysis of nucleotide composition at the junction of translocation events in (**d**).

Polθ expression is essential for the proliferation of HR-defective cells^5,6^, including ones lacking *BRCA1* and *BRCA2*. To examine the function of the different Polθ domains, we treated *Polq*^*ΔHel*^, *Polq*^*ΔPol*^, *Polq*^*Rad51*^, and control *Polq*^+/+^ cells with shRNA targeting *BRCA1*. We then compared the survival of *BRCA1* depleted cells to those treated with control shRNA (Figure 1b and S1b). The results of the colony formation assay indicated that the growth of *BRCA1* depleted *Polq*^*ΔHel*^ and *Polq*^*ΔPol*^ cells was significantly compromised when compared to *Polq*^+/+^ cells (Figure 1b). The findings are in agreement with previous reports that have implicated Polθ helicase and polymerase activities in the survival of cancer cells^5,6,16^. On the other hand, the survival of *Polq*^*ΔRad51*^ cells following *BRCA1* depletion was similar to wildtype cells (Figure 1b). These results suggest that both polymerase and helicase enzymatic activities are important for proper Polθ function, whereas its interaction with Rad51 is dispensable for the survival of HR defective ES cells.

### Polθ–helicase activity promotes efficient repair by alt-NHEJ

We next set out to investigate the involvement of the different Polθ domains during alt-NHEJ. To that end, we tested whether the *Polq* mutations impair chromosomal translocation formation in mES cells, an event that was previously reported to be mediated by alt-NHEJ^36^ and dependent on Polθ^5^. The quantitative chromosomal translocation assay involves simultaneous cleavage of the Rosa26 (chromosome 6) and the H3f3B (chromosome 11) loci with CRISPR/Cas9 (Figure 1c and S1c). Cells transfected with Cas9-gRNA(Rosa26:H3f3b) plasmid were seeded in 96-well plates and a nested PCR was performed to detect breakpoint junctions of chromosomal translocations in each well. We calculated the translocation frequency based on the ratio of positive wells relative to the total number of transfected cells. It is necessary to monitor translocation frequency on the basis of individual wells since population-based PCR methods preferentially amplify shorter transcripts and bias against translocations with larger inserts.

We analyzed translocations in *Polq* mutant cells, and found that cells lacking the Rad51 interaction motif exhibited similar translocation frequencies as wild-type cells (Figure 1d). On the other hand, we observed a significant reduction in the frequency of translocations in *Polq*^*ΔHel*^ as well as *Polq*^*ΔPol*^ cells (Figure 1d). In summary, these data indicate that in addition to the previously established role for the polymerase activity during alt-NHEJ^5,7,16,17^, the Polθhelicase domain is essential for efficient joining of DNA ends in mammalian cells.

To gain additional insight into the function of the different Polθ activities, we examined the sequence of fusion breakpoints at derivative chromosome 6 (Der (6)). We categorized the different translocation events and assigned an alt-NHEJ-signature score based on the presence of insertions and microhomologies (Figure S1d). Interestingly, the alt-NHEJ signature was enriched at translocation events retrieved from wild-type cells but significantly reduced in cells with compromised Polθ-helicase and polymerase functions. Specifically, translocations in *Polq*^*ΔHel*^ and *Polq*^*ΔPol*^ cells exhibited a significant decrease in the frequency of microhomology and nucleotide insertions (>2 nts) (Figure 1e). In contrast, *Polq*^*ΔRad51*^ displayed a similar translocation profile as wild type cells, further confirming that the Rad51 interaction motif is dispensable for Polθ function in alt-NHEJ (Figure 1e and Table S1-9).

### Enhanced HR-driven genome editing by CRISPR/Cas9 upon Polθ inhibition

Having established a role for both Polθ-helicase and polymerase during alt-NHEJ, we next investigated the involvement of these enzymatic activities during suppression of HR, a key phenotype associated with Polθ overexpression and observed in many cancers^5,6^. Using mES cells harboring the different *Polq* mutations, we first measured the formation of IR-induced Rad51 foci, an indirect readout for HR. We detected a significant increase in Rad51 foci in *Polq*^*ΔHel*^ and *Polq*^*ΔPol*^ cells. On the other hand, the accumulation of Rad51 in *Polq*^*ΔRad51*^ cells was similar to wildtype cells (Figure S2a). The latter finding is inconsistent with recent observations implicating the Rapd51 interaction domain in the suppression of HR by polθ^6^. This discrepancy might be attributed to differences in the requirements of Polθ–mediated repair between tumor cells and mouse ES cells. Alternatively, given that the overall levels of Rad51 in cells exceed those of Polθ, it is possible that Polθ can only impact Rad51 function when the polymerase is overexpressed, as is the case in high grade ovarian cancers^6^.

In an independent set of experiments, we aimed to investigate the function of Polθ during HR using a more direct assay. To do so, we investigated whether Polθ inhibition increases HR-mediated gene targeting stimulated by CRISPR/Cas9. We designed a gene targeting assay in which the coding sequence of a Zsgreen fluorescent protein was integrated at the 3’ end of the housekeeping *Hsp90* gene (Figure 2a and S2b-c). Based on this assay, cells that have undergone productive HR are stably marked with green fluorescence and distinguished by fluorescence-activated cell sorting (FACS). Notably, FACS analysis revealed two distinct populations of Zsgreen positive cells (Figure 2b) corresponding to mono-allelic and bi-allelic targeting (Figure 2c). We noticed that targeting of the *Hsp90* locus in MEFs and mES cells was minimally responsive to DNA-PKcs inhibition, suggesting that C-NHEJ is not a major barrier to gene editing at this locus (Figure S2d). Interestingly, we observed a significant increase in the efficiency of *Hsp90* targeting in *Polq*^-/-^ cells relative to *Polq*^*+/+*^ cells (Figure 2d-e and S2d). Furthermore, a similar increase in mono- and bi-allelic targeting of the *Hsp90* locus was detected in *Polq*^*ΔHel*^ and *Polq*^*ΔPol*^ cells (Figure 2e).

**Figure 2.**
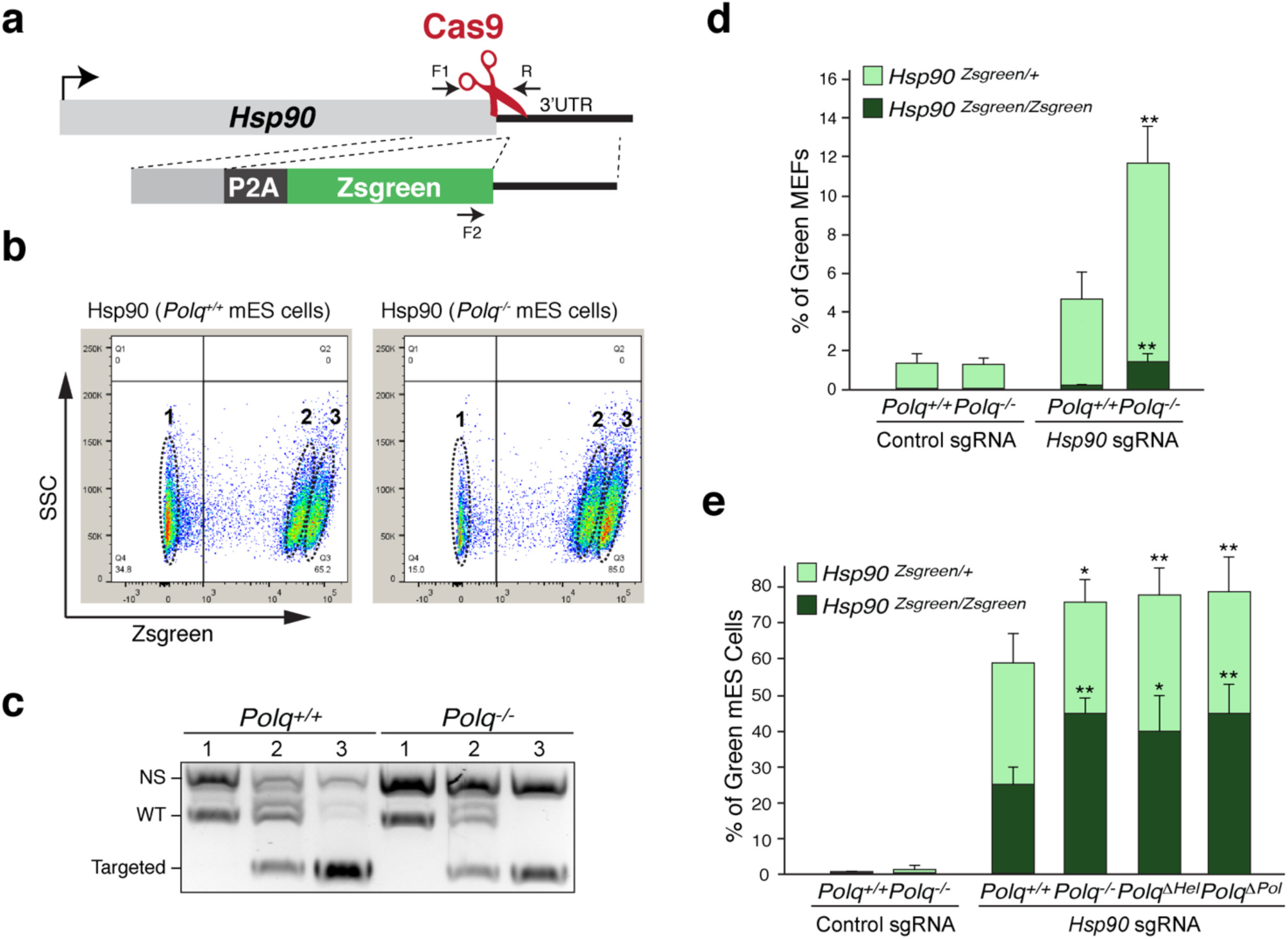
Polθ inhibition increase the efficiency of HR-mediated CRISPR-Cas9 gene targeting. **(a)** Scheme depicting gene targeting assay at the Hsp90 locus. The donor plasmid contains a Zsgreen coding sequence and 600 base pairs of homology arms. (**b**) (FACS) analysis to determine the percentage of ZsGreen positive cells. Three distinct populations were isolated (highlighted as 1, 2 and 3). (**c**) Genotyping PCR for *Hsp90* on DNA corresponding to the three highlighted populations of cells (indicated in (**b**)). (**d**) Gene-targeting efficiency at the *Hsp90* locus in *Polq*^-/-^ and *Polq*^+/+^ MEFs. Bars represent mean ± S.D. (n = 3). (**e**) Gene-targeting efficiency at the *Hsp90* locus in mES cells with the indicated genotype. Bars represent mean ± S.D. (n = 3). **p<0.01; two-tailed Student’s t-test.

We then employed a similar CRISPR/Cas9 strategy to target the highly transcribed *Sox2* gene in mouse ES cells. In contrast to targeting of the *Hsp90* locus, we observed an increase in the percentage of cells expressing green fluorescent protein from the *Sox2* locus in response to DNA-PKcs inhibition, suggesting that this particular locus is more amenable to repair by C-NHEJ (Figure S2e-h). However, depletion of *Polq* did not yield enhanced gene targeting (Figure S2e-h), possibly denoting a minor role for alt-NHEJ in the repair of DSBs incurred at this locus. It was recently reported that the nature of the Cas9-induced lesion influences repair pathway choice, whereby lesions with overhangs preferentially provoke HR and alt-NHEJ over C-NHEJ^37^. We therefore tested the effect of Polθ depletion on HR-mediated repair of breaks triggered using Cas9 nickase (D10A) and a pair of gRNAs that cleave on opposite DNA strands, thereby generating single stranded overhangs. Consistent with previous reports^37^, we noticed a tenfold increase in *Sox2* gene targeting by HR when breaks were induced by Cas9 nickase (Figure S2h-i). In addition, we noticed a small but significant increase in efficiency of gene targeting in *Polq*^-/-^, *Polq*^*ΔHel*^, and *Polq*^*ΔPol*^ cells compared to control cells (Figure S2i). In summary, the data obtained from the gene targeting assay demonstrate that both, helicase and polymerase activities, are essential for suppression of HR by Polθ. In addition, our data suggest that Polθ inhibition could enhance precise genome editing by canonical HR, albeit in locus-specific manner and its effect is influenced by the nature of the ends generated by Cas9 cleavage. Genome-wide-based approaches will be necessary to systematically query the genome and identify specific loci that would yield increased gene targeting in response to Polθ inactivation.

### Polθ-helicase dissociates RPA to promote the annealing of complementary DNA

To investigate the molecular mechanism by which Polθ-helicase, which lacks detectable DNA unwinding activities, promotes alt-NHEJ, we performed a set of *in vitro* experiments. Consistent with previous reports, purified human Polθ-helicase (aa 1-894) displayed robust ATPase activity (Figure S3a-b)^23^. Considering that alt-NHEJ likely requires active annealing between 3’ ssDNA overhangs, we examined whether Polθ-helicase functions as an annealing helicase. For example, helicases such as HARP1/SMARCAL1 have been shown to promote ssDNA annealing in the presence of RPA which acts as a physical and energetic barrier to annealing^38^. Given the previously reported role for replication protein A (RPA) acting as a negative regulator of alt-NHEJ in yeast^31^, we hypothesized that Polθ-helicase utilizes the energy of ATP to overcome the RPA barrier to annealing, in a similar manner to HARP1/SMARCAL1.

To test this hypothesis, we performed ssDNA annealing in the presence of RPA and Polθ-helicase as follows: A 33-nucleotides ssDNA was pre-incubated with RPA in the presence or absence of ATP using standard buffer with MgCl_2_ (Figure 3a). Next, increasing amounts of Polθ-helicase was added for 15 min, followed by the addition of ^32^P-labelled complementary ssDNA for 10 min. Reactions were then terminated and resolved on non-denaturing gels. As previously reported^31,39^, pre-loading of RPA onto ssDNA prevented spontaneous annealing of the complementary DNA substrate (Figure. 3b). Strikingly, the addition of Polθ-helicase to the reaction stimulated DNA annealing in an ATP-dependent manner (Figure 3b). We compared ssDNA annealing reactions with ATP versus a non-hydrolyzable ATP analog (AMP-PNP). The results show that Polθ–helicase annealing occurs in the presence of ATP, but not AMP-PNP, which demonstrates that ATP hydrolysis is required to promote annealing in the presence of RPA (Figure 3b). These data suggest that the helicase enzymatically dissociates RPA from ssDNA in order to promote annealing. Lastly, to address the specificity of Polθ-helicase annealing activity, we performed the annealing reactions using E. coli single-strand binding protein (SSB). In contrast to the results with RPA, Polθ-helicase was unable to promote annealing in the presence of SSB (Figure S3c), which suggests that the helicase was selected to act with RPA during annealing.

**Figure 3.**
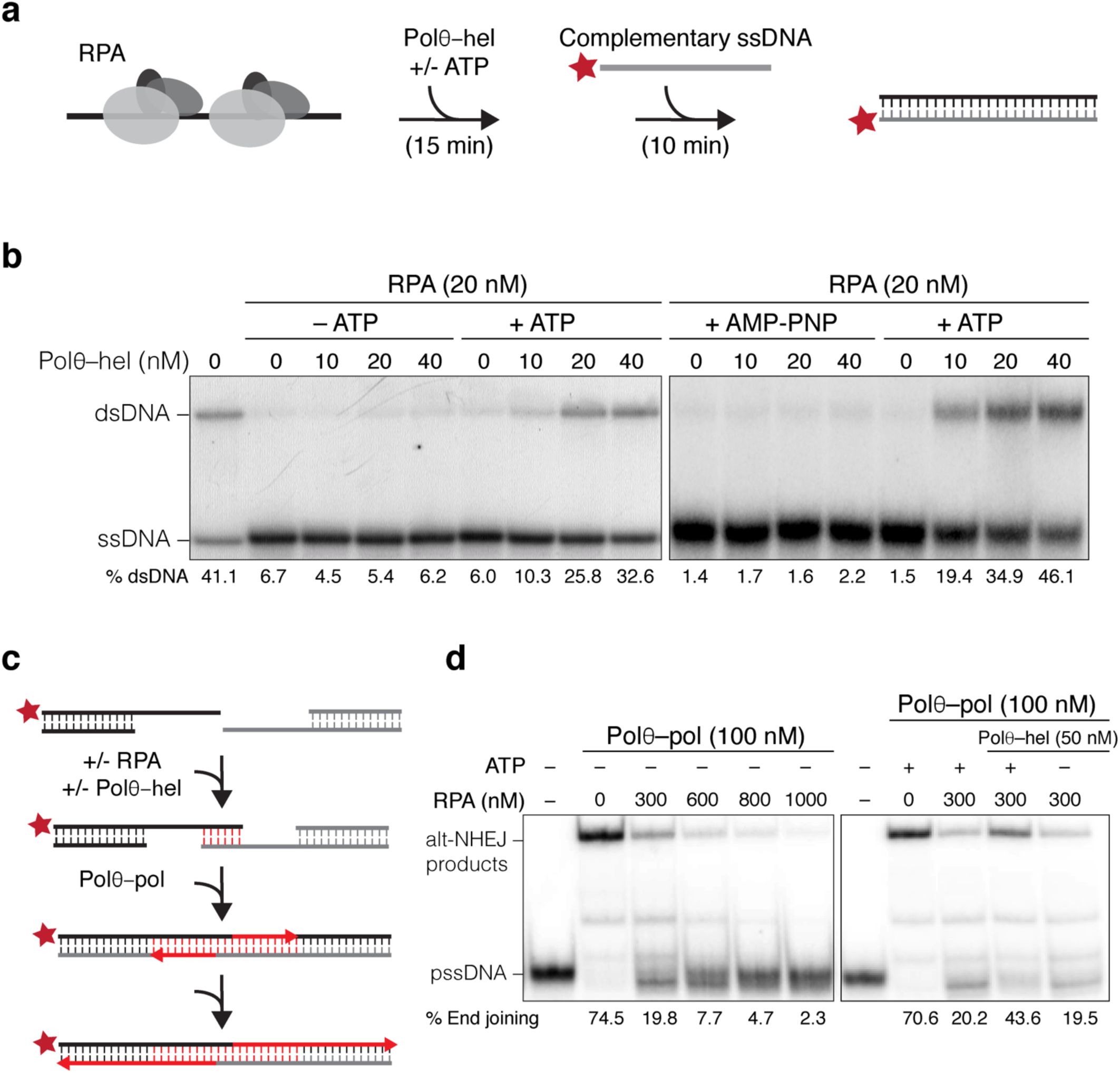
Polθ–helicase antagonizes RPA to promote annealing and alt-NHEJ *in vitro.* **(a)** Diagram of the annealing assay used to investigate whether Polθ–helicase (Polθ–hel) promotes pairing of complementary ssDNA bound by RPA. ssDNA is incubated with RPA, prior to addition of Polθ–helicase. End-labeled complementary ssDNA (indicated with asterisk) is then added, the reaction terminated, and DNA analyzed in non-denaturing gels following protein degradation. (**b**) Representative non-denaturing gels displaying ssDNA annealing in the presence and absence of ATP/AMP-PNP and indicated amounts of Polθ–helicase. % dsDNA indicated. (**c**) Schematic of Polθ–polymerase (Polθ–pol) mediated alt-NHEJ assay. Partially resected DNA model substrates containing 3’ terminal microhomology (6 bases) were used as described^17^. (**d**) Non-denaturing gels showing Polθ–polymerase mediated alt-NHEJ in the presence of the indicated proteins and ATP. % end-joining indicated.

### Polθ-helicase utilizes the energy of ATP to promote alt-NHEJ

The results outlined above implicate Polθ-helicase involvement in the synapsis of resected DSBs and underscore opposing activities exerted by RPA and Polθ-helicase during alt-NHEJ. To gain further insight into the interplay between these two repair factors, we investigated end-joining *in vitro*. To that end, we used partially resected model substrates containing 6 bases of terminal microhomology, previously shown to support efficient end-joining by Polθ–polymerase *in vitro* (Figure 3c)^17^. Consistent with previous results^17^, Polθ–polymerase joined these substrates together forming double-size products in the presence of deoxyribonucleoside triphosphates (dNTPs) as observed in a native gel (Figure 3d). Since RPA suppresses alt-NHEJ in yeast, we tested whether human RPA inhibits end-joining *in vitro*. Indeed, Polθ– polymerase end-joining was significantly reduced when the resected substrates were pre-bound by RPA (Figure 3d and S3d). In contrast, Polθ–polymerase activity was completely resistant to RPA when tested in a traditional primer-template assay (Figure S3e). Taken together, these data demonstrate that the initial ssDNA annealing/synapse step is suppressed by RPA, whereas polymerase extension of the minimally paired overhangs, which is essential for stabilizing the end-joining product, is refractory to RPA binding. To assess whether Polθ–helicase catalytic activity overcomes the RPA block to Polθ–polymerase end-joining, we repeated the reaction in the presence of the helicase domain. Notably, the data demonstrate that the helicase reduces the RPA barrier to end-joining exclusively in the presence of ATP (Figure 3d). The stimulation of DNA annealing by Polθ–helicase was observed under multiple concentrations of RPA and Polθ–helicase (Figure S3d). Altogether, our data indicate that RPA negatively regulates alt-NHEJ by blocking the initial ssDNA annealing/synapse step which is essential for subsequent extension of the minimally paired overhangs by Polθ–polymerase. Notably, Polθ–polymerase alone can promote end-joining in the absence of RPA (Figure 3d and ^17^). In this case, the polymerase presumably takes advantage of transiently and spontaneously annealed/synapsed overhangs which does not require an enzymatic annealing function. However, in the presence of RPA which blocks spontaneous annealing/synapse formation, Polθ–helicase enzymatic activity is needed to reduce this energetic barrier in order to facilitate overhang pairing/synapsis and subsequent Polθ–polymerase extension (Figure 3d and S3d).

### Mammalian RPA inhibits alt-NHEJ *in vivo* and is counteracted by Polθ–helicase

Our *in vitro* data indicated that RPA prevents DNA synapse formation during alt-NHEJ, and that this inhibition is overcome by Polθ–helicase activity to promote end-joining. As mentioned earlier, inhibition of alt-NHEJ by RPA was first detected in *S. cerevisiae*. Specifically, a hypomorphic *rfa1-*D228Y allele that is unable to unwind secondary DNA structures – including ones that resemble synapsed DNA^39,40^ – triggered a 350 fold increase in the frequency of alt-NHEJ^31^. RPA binds ssDNA with similar affinity across eukaryotes and the aspartic acid residue (D228 in budding yeast) is highly conserved (Figure S4a), prompting us to ask whether mammalian RPA (D258 in mice) prevents alt-NHEJ *in vivo*. To that end, we examined DSB repair in the context of dysfunctional telomeres. It has been previously established that upon depletion of the protective shelterin complex from *TRF1^F/F^TRF2^F/F^Lig4^-/-^*Cre-ER^T2^ MEFs, ∼10% of telomeres ends are processed by alt-NHEJ and ∼5% undergo telomere sister chromatid exchange (T-SCE), indicative of HR^41^. As expected, we observed a significant reduction in the frequency of T-SCEs following shelterin loss in MEFs with reduced levels of RPA1 (Figure 4a and Figure S4b-c). In addition, depletion of RPA1 lead to a concomitant increase in alt-NHEJ, denoted by the accumulation of chromosome end-to-end fusions (Figure 4b and S4b-c). We then transduced shelterin-free MEFs with shRNA-resistant RPA1 allele harboring a D258Y mutation, prior to deletion of endogenous RPA1. Analysis of metaphase chromosomes revealed that the expression of mutant RPA failed to rescue the telomere phenotypes associated with RPA1 depletion. In a control experiment, we observed that the expression of a wildtype RPA1 allele restored telomere recombination and prevented the increase in alt-NHEJ (Figure 4a-b and S4a-e). Lastly, we showed that the inhibition of HR through depletion of *Rad51*, which acts downstream of RPA, resulted in a reduction in the frequency of telomere recombination without impacting the levels of alt-NHEJ (Figure S4f-g). In conclusion, these results suggest that RPA acts as a specific regulator of repair pathways choice, promoting error-free repair by HR, while preventing repair by alt-NHEJ.

**Figure 4.**
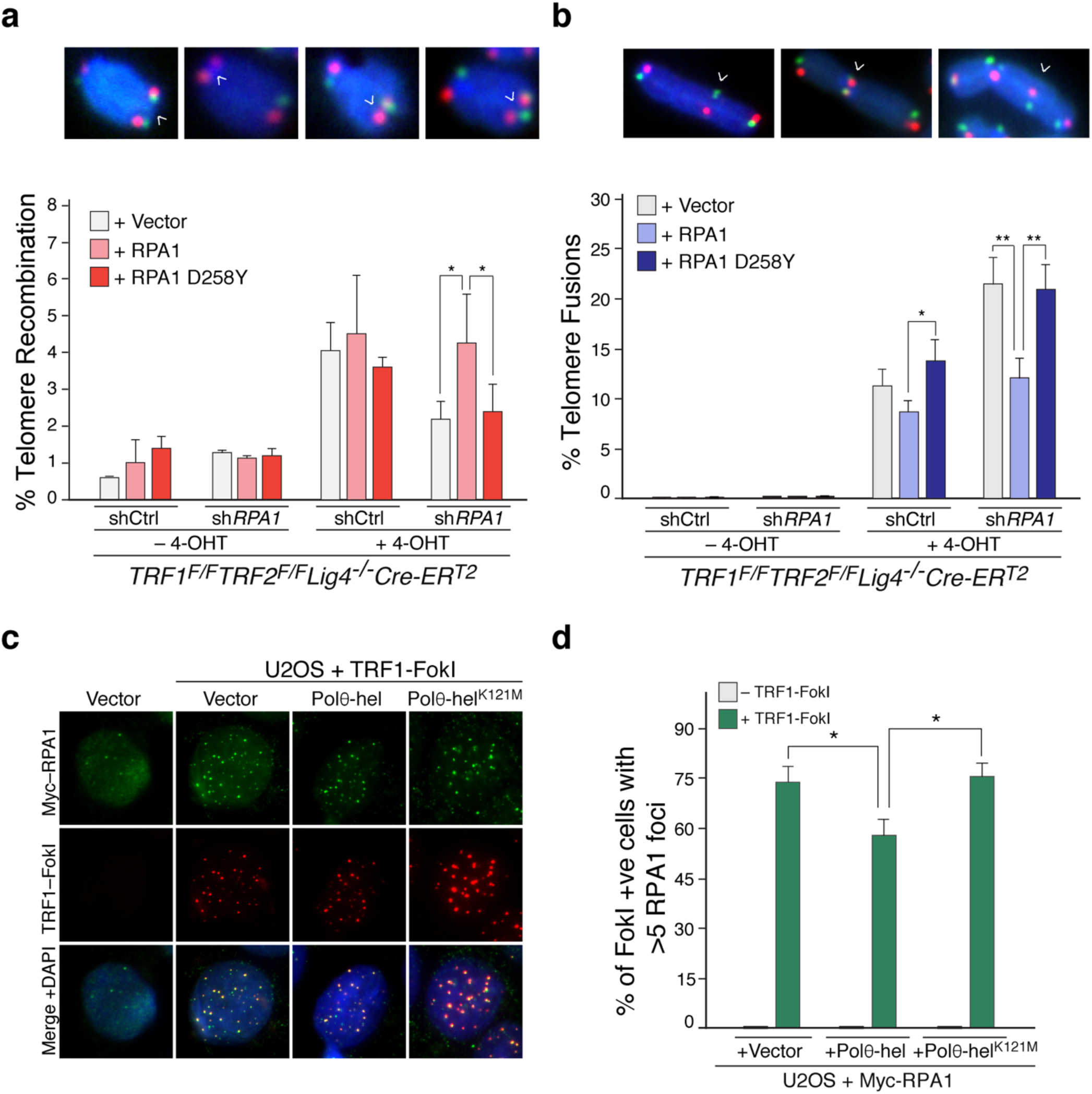
Mammalian RPA inhibits alt-NHEJ. **(a)** Quantification of telomere sister-chromatid exchanges (T-SCEs), reflective of telomere recombination events in *TRF1^F/F^TRF2^F/F^Lig4^-/-^*Cre-ERT^2^ cells with the indicated treatments. Top – Representative examples of T-SCE events (white arrows). Error bars denote S.E.M. from two independent experiments. *P < 0.05, **P < 0.01; two-tailed Student’s t-test. **(b)** Quantification of telomere fusions by alt-NHEJ in cells with the indicated treatment. Top – Examples of chromosomes fusion events denoted by white arrows. Bars represent mean ± S.D. from three independent experiments. *P < 0.05 and **P < 0.01; two-tailed Student’s t-test. (**c**) Immunofluorescence for Myc-RPA1 and TRF1-FokI-mCherry in U2OS cells expressing the indicated alleles. TRF1-Fok1-mCherry expression was induced upon treatment with doxycycline, shield-1, and 4-OHT. (**d**) Graph representing the quantification of Myc-RPA1 accumulation in cells expressing TRF1-FokI (as in c). Bars represent mean ± S.D. from three independent experiments. *P < 0.05 and **P < 0.01; two-tailed Student’s t-test.

Finally, we sought to examine the effect of Polθ–helicase on the accumulation of RPA at DSBs *in vivo*. In order to visualize RPA at break sites, we turned to the previously established TRF1-Fok1 system that triggers multiple DSBs per telomeres^42^. It was recently noted that Fok1-induced telomere breaks were not processed by C-NHEJ, but fixed by HR and alt-NHEJ^43^, which renders them ideal substrates to examine the interplay between RPA and Polθ–helicase *in vivo*. The expression of TRF1-FokI fusion protein in U2OS cells lead to significant co-localization of Myc-RPA1 at damaged telomeres (Figure 4c). Interestingly, we observed a small, but significant reduction in the accumulation of RPA1 at telomere breaks in cells expressing Polθ–helicase (Figure 4c and Figure S4h-i). Importantly, expression of a helicase-defective Polθ allele (K121M) did not impact the formation of RPA1 foci at telomere breaks (Figure 4c and Figure S4h-i). Taken together, these results highlight a role for Polθ–helicase in displacing RPA from ssDNA while catalyzing overhang annealing, thereby facilitating alt-NHEJ at the expense of HR.

## Discussion

Error-prone repair by alt-NHEJ can destabilize the genome due to the accumulation of insertions and deletions at break sites. Alternatively, repair of similar lesions by HR will lead to a safer outcome. Paradoxically, both pathways display maximal activity in S and G2 and share the initial resection step. The underlying mechanism(s) that regulate the choice between HR and alt-NHEJ following the resection is poorly understood. In this study, we provide evidence in support of an antagonistic interplay between Polθ and RPA that orchestrates DSB repair. Specifically, our results highlight a crucial role for the helicase domain within Polθ to promote alt-NHEJ during chromosomal translocation in ES cells, and suppress HR mediated gene editing. In addition, our data establish that Polθ–helicase offsets the inhibitory role of RPA to foster the annealing of microhomologous sequences between opposing ssDNA overhangs generated at DSBs. Finally, we show that inhibition of RPA, or expression of an RPA-D258Y mutant that is unable to block spontaneous ssDNA annealing, leads to significant inhibition of HR and a concomitant enhancement in alt-NHEJ.

Based on our observations, we propose a model in which a key commitment step to repairing breaks by HR versus alt-NHEJ is executed at the level of DNA end synapsis (Figure. 5). The first decision point during DSB repair is achieved through 5’-3’ DNA resection by MRN-CtIP to block C-NEHJ^3,44,45^. The binding of RPA to the resulting short ssDNA overhangs stimulates further resection by BLM/EXO1 to facilitate subsequent steps during HR^46,47^. In addition, RPA inhibits the spontaneous annealing of microhomologous sequences exposed by end resection^31,39,48^. On the other hand, Polθ–helicase actively dissociates RPA to promote DNA end-synapsis; a critical commitment step for alt-NHEJ (Figures 3 and 4). In effect, the antagonistic activities between Polθ–helicase and RPA define a second decision point during DSB repair immediately following initial end-resection. When Polθ–helicase prevails, the minimally paired ssDNA overhangs are subsequently extended by Polθ–polymerase which is essential for stabilizing the alt-NHEJ intermediate and filling in the gaps prior to covalent joining of DNA by Lig3, or Lig1 (to a lesser extent)^49^.

**Figure 5.**
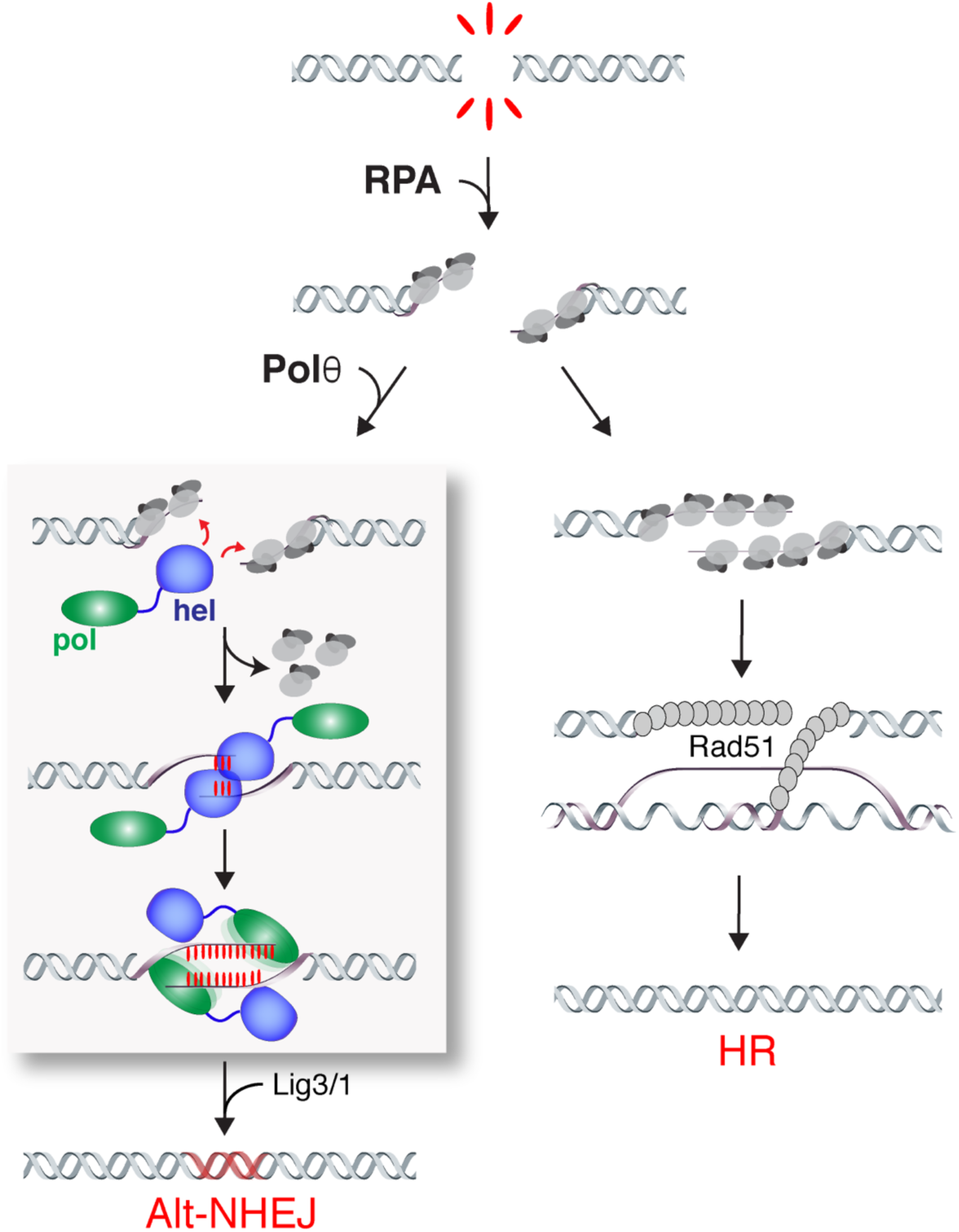
The interplay between Polθ and RPA determine the outcome of DSB repair. Schematic depicting our model for the interplay between Polθ and RPA during DSB repair. Binding of the RPA complex to resected DSB ends block alt-NHEJ and promote HR. The helicase activity of Polθ perturbs the binding of RPA to ssDNA and stimulate the annealing of resected DSBs. Annealed intermediates are then stabilized upon fill-in synthesis by Polθ–polymerase and ultimately joined by Lig3/Lig1.

The helicase domain of Polθ is most closely related to HelQ, also known as Hel308, an ATP-dependent helicase that unwinds replication fork DNA substrates to mediate replication-coupled DNA repair in response to ICL inducing agents^50,51^. In contrast, Polθ is thought to lack DNA unwinding activity^23,24,52^ and mammalian cells lacking *POLQ* are not sensitive to ICL^16,33^, thus highlighting significant functional divergence between the two helicases. The annealing function of Polθ–helicase is highly reminiscent to that displayed by HARP/SMARCAL1, an annealing helicase that is recruited to sites of replication stress *via* an interaction with RPA to minimize the amount of ssDNA and protect stalled replication forks^38,53,54^. We were unable to detect a direct interaction between RPA and Polθ (data not shown), suggesting that the recruitment of Polθ to DSBs is independent of the RPA complex. Whether Polθ utilizes its annealing activity to allow cancer cells to better tolerate replication stress^55^ remains to be addressed.

RPA is a major eukaryotic ssDNA binding protein that has been shown to unwind secondary structure and antagonize annealing of complementary oligonucleotides^31^. Consistent with this activity, yeast strains carrying RPA1 mutations exhibited enhanced alt-NHEJ activity^31^. Our results demonstrate that the inhibitory role for RPA during alt-NHEJ is conserved in higher eukarotes. Nonetheless, there remain significant differences between alt-NHEJ in yeast and mammals. One difference is illustrated by the degree of microhomoloqy required for joining broken DNA. Alt-NHEJ events in budding yeast requires >5 bp of microhomoloqy and are less tolerant to mismatches^28,29,30^, whereas joining in mammals can take place with as little as one nucleotide of homology^32^. The thermostability of base pairing might be sufficient to promote end-joining in yeast. Alternatively, metazoans acquired an essential and dedicated activity – that of Polθ – to enzymatically promote annealing/synapse of ssDNA overhangs with limited microhomoloqy and rapidly stabilize these minimally paired ends via their extension by the polymerase domain. Given that yeast lack a *Polq* gene, it will be interesting to test whether the expression of Polθ alleviates the requirement for extensive microhomoloqy during alt-NHEJ-mediated repair in S. *cerevisiae*.

*POLQ* is frequently overexpressed in human cancers^11,56^, and its overexpression is linked to poor prognosis in breast cancer^34^. Furthermore, *POLQ* expression confers resistance to DSB forming agents, including ionizing irradiation and chemotherapy drugs^16,18^. Lastly, Polθ–mediated alt-NHEJ was proposed to be an adaptive mechanism that allows the survival of cells with defective HR or C-NHEJ, including *BRCA1*/2 mutated breast and ovarian cancer cells^5-7^. As a result, Polθ has emerged as a novel cancer drug target. Our findings that both enzymatic activities support Polθ function during DSB repair highlight more opportunities for targeting this unique enzyme in cancer.

## Authors contribution

Experiments were designed by A.S., R.T.P., P.A. M-G. and T.K. T.K. performed in vitro experiments. T.K. and E.K. purified proteins. P.A.M-G. performed all other experiments. A.S. wrote the manuscript. All authors discussed the results and commented on the manuscript.

### Acknowledgements

We thank R. Greenberg and T. de Lange for providing key reagents. We are grateful to Sarah Deng, Alexandra Pinzaru, and Raymond Barry for providing comments on the manuscript. This work was supported by grants from Pershing Square Sohn cancer research alliance (A.S.), the V-foundation (A.S.), Pew-Stewart scholars award (A.S.), and the National Institutes of Health award 1R01GM115472-01 (R.T.P.). P.A. M-G. is supported by a fellowship from the Department of Defense (BC134020).

